# Predicting regional somatic mutation rates using DNA motifs

**DOI:** 10.1101/2022.08.04.502732

**Authors:** Cong Liu, Zengmiao Wang, Jun Wang, Chengyu Liu, Mengchi Wang, Vu Ngo, Wei Wang

## Abstract

How the locus-specificity of epigenetic modifications is regulated remains an unanswered question. A contributing mechanism is that epigenetic enzymes are recruited to specific loci by DNA binding factors recognizing particular sequence motifs (referred to as epi-motifs). Using these motifs to predict biological outputs depending on local epigenetic state such as somatic mutation rates would confirm their functionality. Here, we used DNA motifs including known TF motifs and epi-motifs as a surrogate of epigenetic signals to predict somatic mutation rates in 13 cancers at an average 23kbp resolution. We implemented an interpretable neural network model, called contextual regression, to successfully learn the universal relationship between mutations and DNA motifs, and uncovered motifs that are most impactful on the regional mutation rates such as TP53 and epi-motifs associated with H3K9me3. Furthermore, we identified genomic regions with significantly higher mutation rates than the expected values in each individual tumor and demonstrated that such cancer-specific regions can accurately predict cancer types. (The code is available from https://github.com/Wang-lab-UCSD/SomaticMutation)

**Significance Statement:** The relationship between DNA motifs and somatic mutation rates in cancers is not fully understood, especially at high resolution. Here we developed an interpretable neural network model to successfully predict somatic mutation rates using DNA motifs in 13 diverse cancers and identified the most informative motifs. Furthermore, we showed that the genomic regions with significant higher mutation rates than the predicted values can be used for cancer classification.

## Introduction

Locus-specific epigenetic modifications, including DNA methylation and histone modifications, play critical roles in development and other key biological processes (1). While epigenetic patterns are affected by nucleosome positioning (2), modifying enzymes (3), transcription factors (TFs) (4), non-coding RNAs (ncRNAs) (5), signaling molecules (6) and three-dimensional genomic organization (7, 8), the epigenetic modifying enzymes generally do not recognize specific DNA sequence or do not bind to DNA at all and they need to be recruited to specific loci by DNA binding proteins or ncRNAs. Pioneer transcription factors are examples of such proteins that bind to the condensed chromatin to initiate chromatin remodeling and activation of regulatory elements in particular loci (9–12). However, proteins and their binding motifs responsible for establishing or maintaining other types of locus-specific epigenetic patterns largely remain elusive.

Accumulating evidence starts to reveal the importance of DNA sequence features in shaping epigenetic patterns (4, 13–20) and DNA motifs associated with epigenetic modifications (referred to as **epi-motifs**) have also been documented (21–23). The readout of these epi-motifs is dynamic and dependent upon cellular conditions (e.g. activity of the DNA binding regulator and its access to DNA), and thus is the epigenome. This mechanism is similar to how TFs function: while the TF motifs remain the same, the transcriptional regulation is tissue-specific and dynamic. As successful prediction of gene expression using TF motifs supports the functionality of TF motifs (24, 25), we reason that using epi-motifs to predict biological outputs depending on local epigenetic state such as somatic mutation rates would help to illustrate their importance in regulating epigenetic locus-specificity.

Somatic mutations are tightly associated with disease phenotypes and are resulted from the interplay between mutagenic processes and DNA repair mechanisms (26–32). The regional mutation rates are related to various factors, including replication timing, transcriptional activity, nucleosome positioning, chromatin accessibility, histone modifications and protein binding (26–32). Analysis of mutation rates has been performed at multiple scales. At the megabase-scale, high mutation rate is correlated with later replication timing, closed chromatin, strong repressive (e.g. H3K9me3) and weak active (H3K4me1/2) histone marks (33–38). At the gene scale, reduced mutation rate is associated with high transcription and high H3K36me3 levels (39–42). At the scale of tens to hundreds of bases, nucleosome positioning is correlated with periodicity of mutation rates (43–49); furthermore, while high mutation rates are observed at the binding sites of CTCF (40, 50, 51), ETS family and numerous other transcription factors (TFs) (52– 55), simultaneous analysis of DNA damage and repair suggested that the impact of protein binding varies from no effect to inhibition or stimulation on DNA damage depending on TF and DNA damaging agent (56). At the smallest scale of several base pairs, previous analyses have uncovered sequence context of somatic mutations and mutational signatures (57–62).

These observations support that chromatin state is tightly correlated with regional mutation rate (29–31). Such a relationship can be quantified at the megabase scale by machine learning models (34, 37, 63). However, no strong correlation between individual epigenetic signals and mutation rates could be observed at finer (e.g. tens of bases to kilobase) scales (34, 37, 63) and no quantitative model has been established, which hinders revealing the epigenetic mechanisms regulating somatic mutation. As protein binding has been suggested to impact the balance between DNA damage and DNA repair rates around their binding sites (52–56), considering DNA motifs recognized by DNA binding proteins may help to establish a prediction model but it has to overcome the challenge that proteins can have divergent effects on mutation rate upon binding (56).

Given that driver mutations in cancers and other diseases only account for a small portion of all somatic mutations, we reason that (1) the majority of somatic mutations are decided by the regional features, such as epigenetic state and TF motifs, and the regional mutation rates can be predicted by these relevant features; in other words, somatic mutations in these regions are disease-independent and their occurrence is only related to the local environment rather than the disease state. (2) a relatively small portion of genomic regions contain disease-specific mutations and the mutation rates in these regions significantly deviate from the expected values; in other words, these mutations are driven by the disease state and have higher mutation loads than expected from the local environment.

We present here an interpretable deep neural network model that predicts somatic mutation rate at the kilobase scale using DNA motifs. We set to calculate the mutation rate in genomic regions with distinct epigenetic states as annotated by ChromHMM (3) based on the following rationale. As the epi-motifs are associated with the regional epigenetic state and the regional epigenetic state has been shown to be associated with somatic mutation rates, it is reasonable to use epi-motifs together with other motifs, which are often associated with DNA damage and repair such as TP53, to predict regional mutation rates in the same epigenetic state. If randomly segmenting the genome, a single region likely contains segments in different epigenetic states (e.g. the active enhancers are often 200-300bp long which is only part of a kilobase-long region) and it is thus irrational to predict the mutation rate of the entire region using epi-motifs (together with other motifs). If randomly segmenting the genome at a high resolution such as 200bp to avoid this problem, the regions is too short to get stable mutation rates and we are not aware of any study that can predict mutation rates at even kilobase resolution only using sequence.

We chose DNA motifs as input features because protein binding has been shown associated with the regional mutation rate (29–31). By building such a predictive model, we aimed to uncover the DNA motifs that enhance or repress somatic mutations (Figure 1). Previously, identification of TF motifs important for regulating gene expression in promoters and enhancers helped to understand transcriptional regulation (64, 65). Similarly, we propose that identification of mutation-associated motifs particularly epi-motifs will help pave the way towards revealing the molecular mechanisms influencing the rate of regional somatic mechanisms.

**Figure 1.**
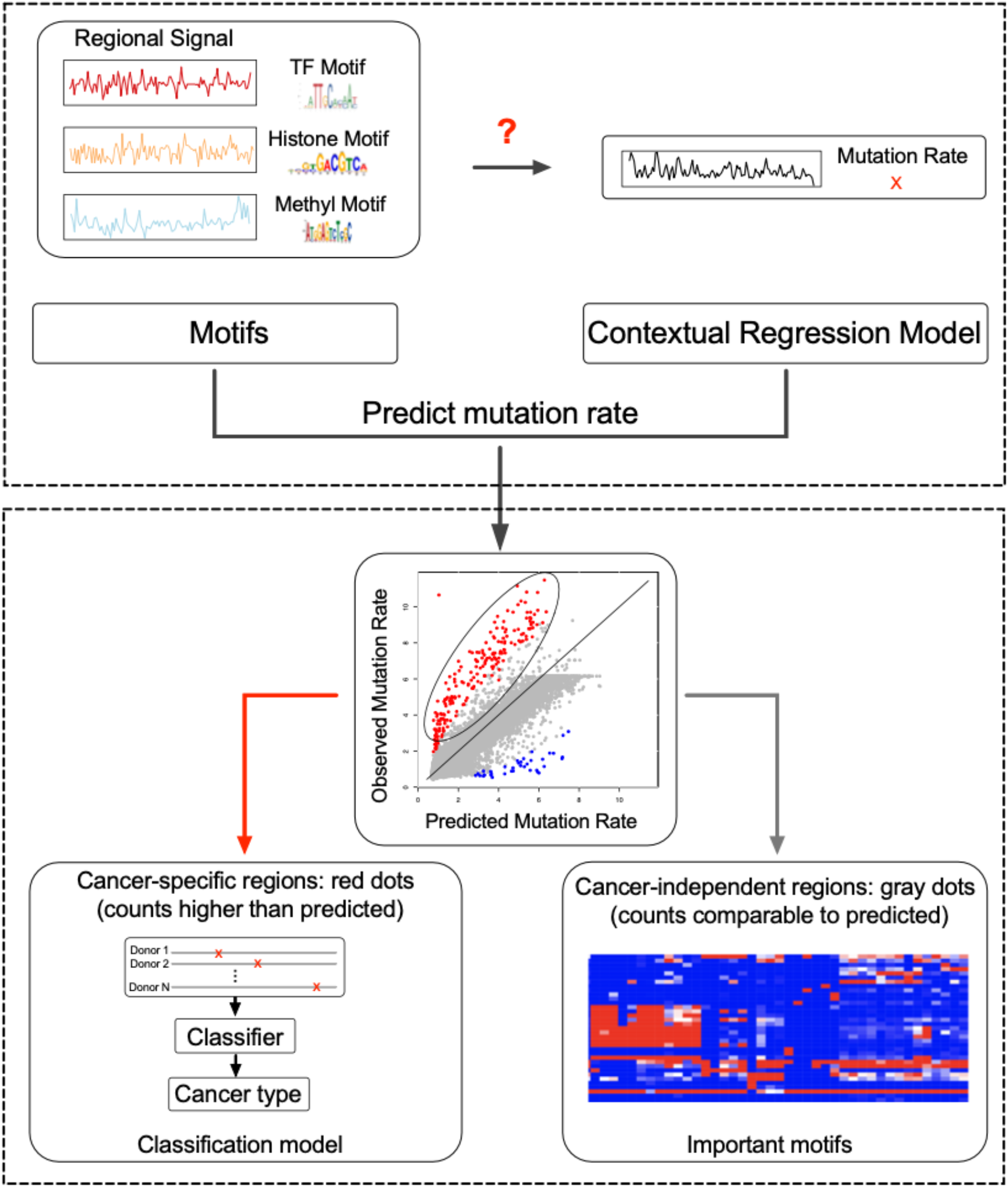
The flowchart of the analysis. Using DNA motifs, including known TF motifs (TF motifs), histone associated motifs (Histone motifs) and DNA methylation associated motifs (Methyl motifs) to represent epigenetic states, we built a contextual regression (CR) model to predict regional mutation rates. As the majority of the mutations are related to the local epigenetic state and independent from the disease state (grey dots), this CR model can quantify the relationship between DNA motifs and somatic mutation rates. Importantly, the CR model revealed the motifs most predictive of somatic mutations (right branch) and the predicted mutation values allowed classification of cancer types using the cancer-specific regions with significantly higher mutation rates than predicted (left branch). In the scatter plot, each point represents a training/testing instance, which is the predicted/measured mutation rate of a genomic region. The mutation rate is the log2(MutationRate+1), which is consistent with Fig. 2c. The rows of the heatmap are important motifs and the columns are different types of cancers.

The model has the following unique features. First, we included not only the known motifs documented in the literature but also de novo motifs that are associated with DNA methylation and histone modifications (referred to as **epi-motifs**) (21–23). Inclusion of epi-motifs can approximate the epigenomic signals and would reveal DNA motifs involved in regulating locus-specific epigenetic modifications. Second, this is an interpretable model: using a method called contextual regression (66, 67), it can assess the contribution of each motif to the prediction accuracy and determine whether the presence of a motif is associated with increase or decrease of regional mutation rate. Third, this model can identify disease-specific regions that contain exceeding loads of mutations and mutations in these regions can be used for classifying disease type.

## Results

### Regional somatic mutation rates could be predicted using DNA motifs

We collected somatic mutations in 1,125 donors detected by whole-genome sequencing (WGS) from the Pan-Cancer Analysis of Whole Genomes (PCAWG) project (68) (see Online Methods for selection criteria and Supplementary Table 1). These donors were from 13 tumor types and contained 8,086,632 somatic mutations. In the normal tissues corresponding to each cancer type, we took the genomic segmentation defined by ENCODE using ChromHMM (69). The logic of this approach is that the majority of the mutations in cancers are random ones and we reasoned that they only depend on the local epigenetic state, which can be approximated using the normal cells. In fact, 80.3% of the genome (on average) have the similar ChromHMM states in the cancer cell line and the corresponding normal cell line (Supplementary Table 2). Because the ChromHMM segments vary in length reflecting the distinct scales of different chromatin states, we calculated and normalized mutation rate in each region as 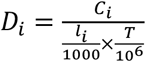, where *T* is the total number of somatic mutations in the tumor dataset under consideration, *C*_*i*_ the number of somatic mutations in region *i, l*_*i*_ is the length (bp) of this region. Note that the region size is much smaller (on average 22.7kb, Supplementary Table 3) than 1Mbp in the previous studies (34, 37, 63). This finer scale is expected to better capture the regional epigenetic state and mutation rate.

To build a prediction model, we considered DNA motifs as input features. We included 1,663 known human motifs documented in the literature as well as 310 and 348 motifs previously identified to be associated with DNA methylation (23) and histone modifications (22), respectively (Online Methods and Supplementary Table 4). We trained and tested a contextual regression (CR) model to predict the regional mutation rates on 80% of all the donors, while the remaining 20% of donors were left out for evaluating the model’s capability for classifying cancer types. The framework of this study is shown in Supplementary Figure 1.

Contextual regression (CR) is a framework to interpret machine learning models (66). It has been successfully applied to identify important features from neural network models such as those predictive of open chromatin (66) and circular RNA biogenesis (67). Here, we constructed a CR model with a fully connected neural network (Figure 2a), including one input layer and 7 hidden layers. The 1st, 3rd and 5th hidden layers respectively contain *p*/2, *p*/10-20, *p*/5-30 nodes (*p* is the number of features) with rectified linear unit (ReLU) activation function, and each is followed by a dropout layer (2nd, 4th, 6th hidden layer) with a rate of 0.01, 0.01 and 0.1, respectively. Dropout layer is a common technique used in model training to prevent model overfitting. Dropout rate refers to the percentage of the neurons that are randomly dropped out in the hidden layer. If the dropout rate is high, it means more neurons are dumped. The 7th layer is the Context Weight layer with *p* nodes and a linear activation function. Dot product between the input and Context Weight layers generates the output (Figure 2a).

**Figure 2.**
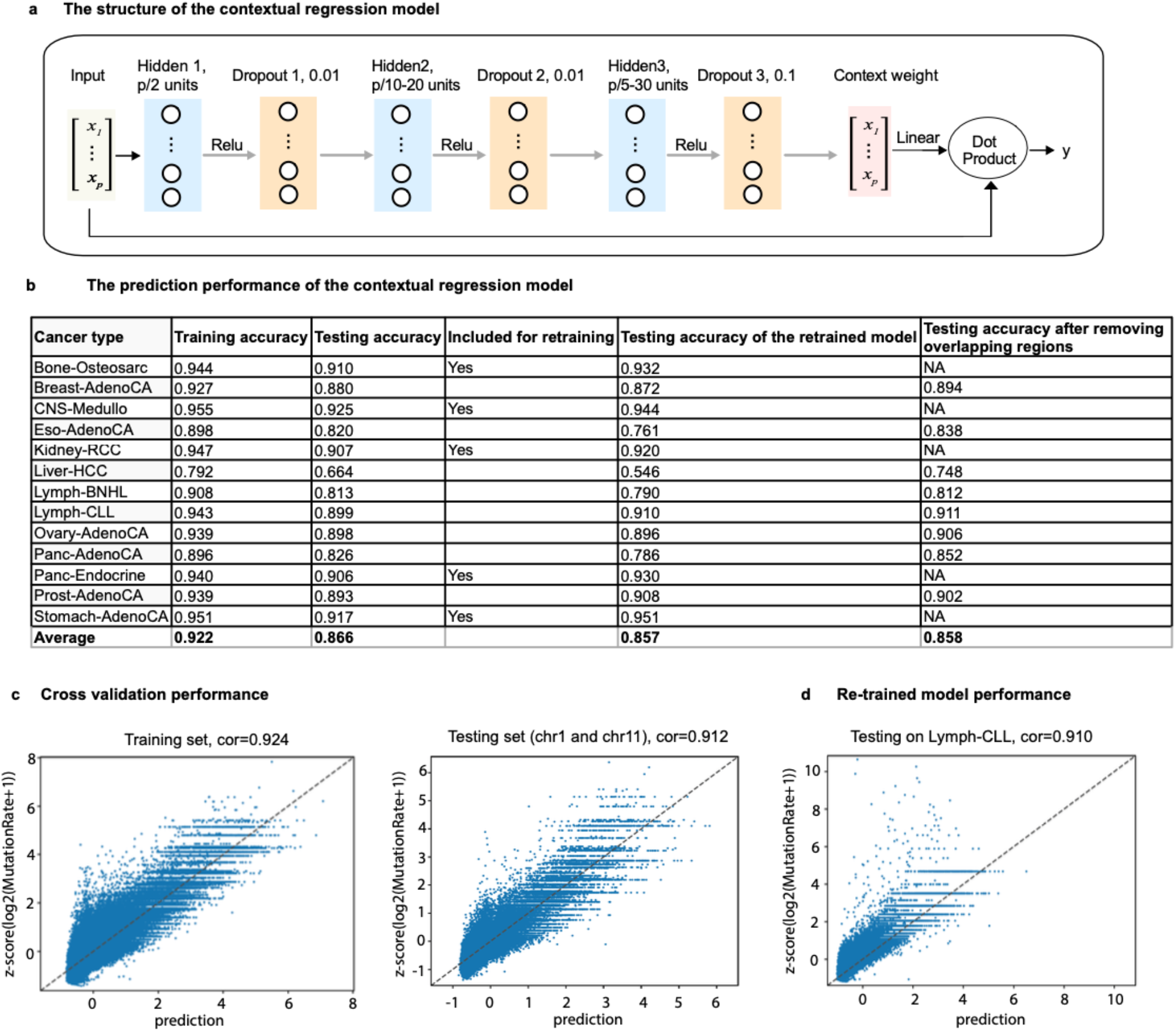
The contextual regression model successfully predicted somatic mutation rates in 13 tumors. (**a**) The structure of the contextual regression model; (**b**) For each tumor type, 10-fold cross validation was performed and the Pearson correlation coefficient was calculated between the predicted and measured values. “Training accuracy” and “Testing accuracy” represent the average of Pearson correlation coefficients in the training, testing datasets respectively. “Included for re-training” indicates which data set was included for re-training the CR model after removing the cancer-specific regions (i.e. the regions with mutation rates significantly deviating from the predicted values). “Testing accuracy of the re-trained model” represents the correlation using the re-trained CR model obtained from the interactive procedure (see Online Methods). Because the regions from a tumor type in the test set may overlap with the regions included in the merged dataset for training, the overlapped regions were removed and Pearson correlation coefficients are shown as “Testing accuracy after removing overlapping regions”; (**c**) The scatter plot for one fold from the 10-fold cross validations in which chr1 and chr11 were left out as the testing set; (**d**) The scatter plot for prediction in Lymph-CLL testing set using the re-trained CR model.

We hypothesized that (1) the majority of the somatic mutations are random and independent of cancer (the associated regions are referred to as **cancer-independent regions**), and (2) such random mutations are associated with the local epigenetic state and can be predicted by DNA motifs. To test this hypothesis, we trained a model to predict mutation rates in all regions and proposed that an accurate prediction would confirm the dominant majority of the regions to be cancer independent. We then identified **cancer-specific regions** as those whose observed mutation rates significantly deviate from the predicted values and the prediction model could be further improved by removing the identified cancer-specific regions (see Supplementary Figure 2).

To implement this strategy, we first performed the CR prediction with 10-fold cross validations in each cancer type and calculated the Pearson correlation between the predicted and observed mutation rates (see Supplementary Table 3). The average Pearson correlation was high (0.866, Figure 2 b), indicating a dominant majority of cancer-independent regions. To build a universal model for all cancers, we selected 5 cancers with large sample size (donors and regions) and superior prediction performance (>0.90 average Pearson correlation coefficients in the test sets in the 10-fold cross validations), including Bone-Osteosarc, CNS-Medullo, Kidney-RCC, Panc-Endocrine and Stomach-AdenoCA. We merged the regions of these five cancers. To avoid the similar regions present in both training and test sets, we held out 2 or 3 chromosomes for testing while training the model on the other chromosomes. We performed 10 such cross-validations and the average Pearson correlations were 0.926 on the training and 0.907 on the test sets, respectively (see Supplementary Table 5 and Figure 2 c). Note that the ChromHMM regions are different for different cancers. By checking the unique ChromHMM states across 13 cancers, we did not find many exact ChromHMM segments, which have exactly the same starting and ending locations in genome, across 13 cancers. Specifically, 80% of the ChromHMM states occur in only one cancer while a negligible portion (much smaller than 0.01%) occurs in all 13 cancers. Therefore, the features on the regions are different for different cancers and their somatic mutation rates can be predicted using the same model.

We next focused on the cancer-independent regions to improve the prediction performance. For this purpose, we first identified and removed regions whose observed mutation rates significantly deviated from the predicted values (Online Methods). Then, we re-trained the CR model using the remaining regions (i.e. cancer-independent regions) that better captured the relationship between epigenetic state and random somatic mutations (Supplementary Figure 2). We next refined the identification of cancer-independent regions and cancer-specific regions using the re-trained CR model. Consistent with our hypothesis, the majority of regions are cancer-independent, ranging from the highest of 90.1% in Breast-AdenoCA to the lowest of 63.1% in Prost-AdenoCA (Figure 3 a, Supplementary Table 6).

**Figure 3.**
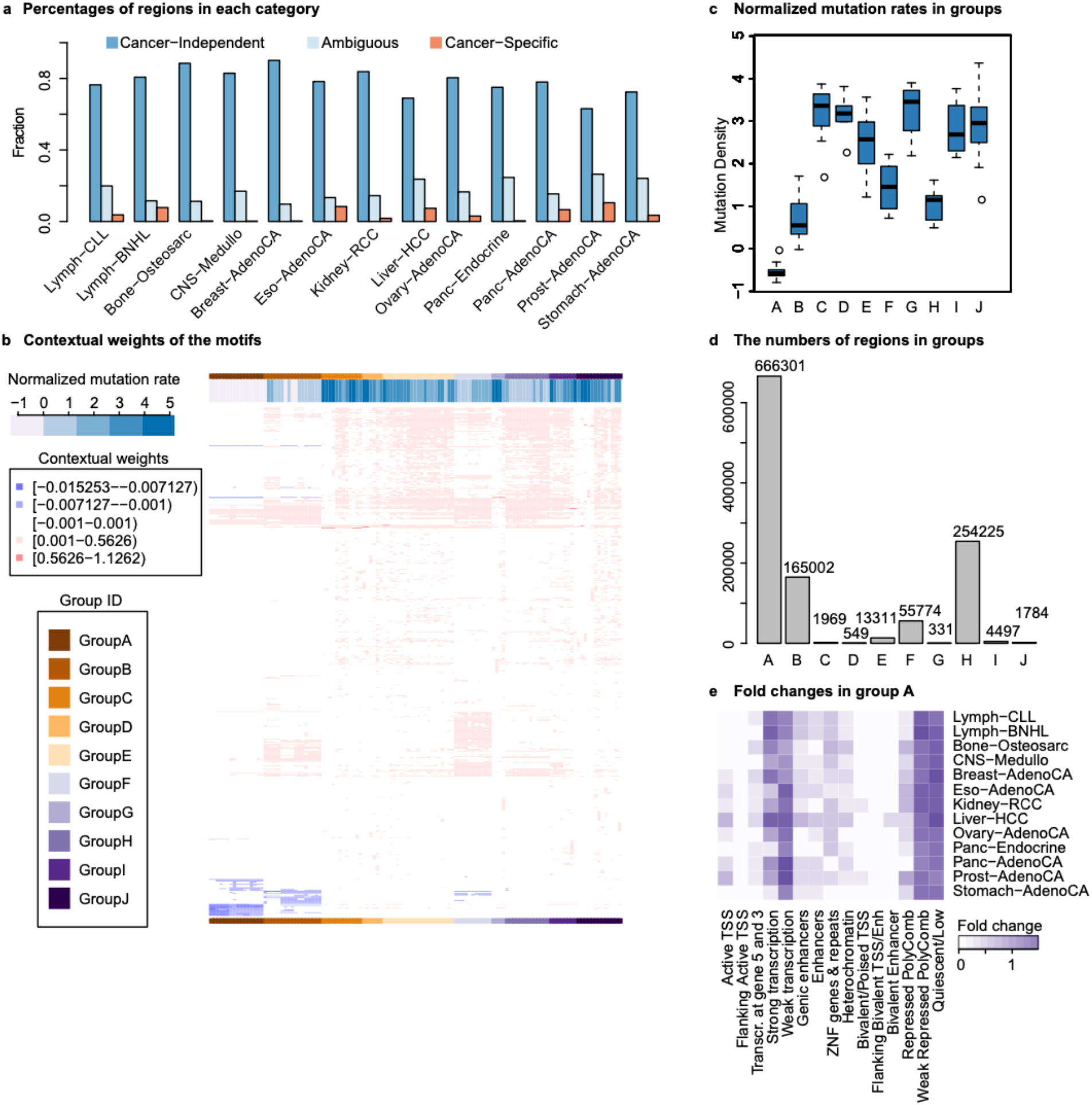
Analysis of the cancer-independent regions. (**a**) The percentage of cancer-independent regions in the 13 cancer types. Percentage is calculated as the number of cancer-independent/specific or ambiguous regions divided by the total number of regions in a cancer; (**b**) Cancer-independent regions clustered using the contextual weights of the motifs. For each of the 13 cancer types, the identified cancer-independent regions were clustered into 10 clusters using the Manhattan distance between the feature contextual weight vectors as the similarity metric. Each row is a motif with non-zero contextual weight, each column a cluster, and each entry is the average of a motif’s contextual weights in all the regions in a cluster. The clusters were further clustered into 10 groups; (**c**) The normalized mutation rate of each group, which is the z-score of mutation density (see methods for more details), varies significantly from the lowest in group A to the highest in group G; (**d**) The numbers of regions in the 10 groups; (**e**) The fold change of chromHMM states in group A and each tumor. The fold change for each chromHMM state is defined as the percentage of the state in group A divided by the percentage of the state in all the regions in a specific cancer.

To evaluate the prediction performance using an independent data set, we applied the re-trained CR model to the remaining 8 cancer types (Figure 2 b and d, Supplementary Table 3, Supplementary Figure 3) and achieved an average Pearson correlation coefficient of 0.857. Considering regions included in the 5 training datasets might also appear in the other datasets, we removed all the overlapping regions in the test set and the average Pearson correlation remained high (0.858, Figure 2 b). Taken together, the CR model successfully captured the relationship between cancer-independent somatic mutation rates and DNA motifs in diverse tissues.

It is worth noting that the prediction performance using DNA motifs alone is comparable to that using ChIP-seq data of TFs and histone modifications. For example, using 165 TF and histone ChIP-seq of data in GM12878 as the input features to predict the somatic mutation rates of Lymph-CLL (i.e. GM12878 as the corresponding normal cell for Lymph-CLL), the Pearson correlations on the training and testing datasets were 0.903 (compared to 0.943 using motifs) and 0.871 (compared to 0.899 using motifs), respectively. The slightly lower correlation using ChIP-seq data may be due to the smaller number of TFs measured by ChIP-seq compared to the number of available motifs. As most cancers do not have many ChIP-seq data in the corresponding normal tissues/cell lines, this observation indicates that using TF and epi-motifs can be useful for approximating the regional epigenetic states in predicting mutation rates.

As the regional mutation rates are associated with the epigenetic state, we analyzed the prediction results to ensure that our model did not only predict the average mutation rates in different ChromHMM states. First, we showed that the same ChromHMM state has a wide range of mutation rates. Supplementary Figure 4 shows the log2(mutation rate) distribution in breast cancer as an example. All the other cancers have the same conclusion. Second, we showed high correlation between the predicted and measured mutation rates in all the regions for the same ChromHMM states (breast cancer as an example in the Supplementary Figure 5 and each panel is one ChromHMM state). These observations clearly showed that our model can indeed predict mutation rates for individual regions and not just distinguish the mean mutation rates between different ChromHMM states.

**Figure 4.**
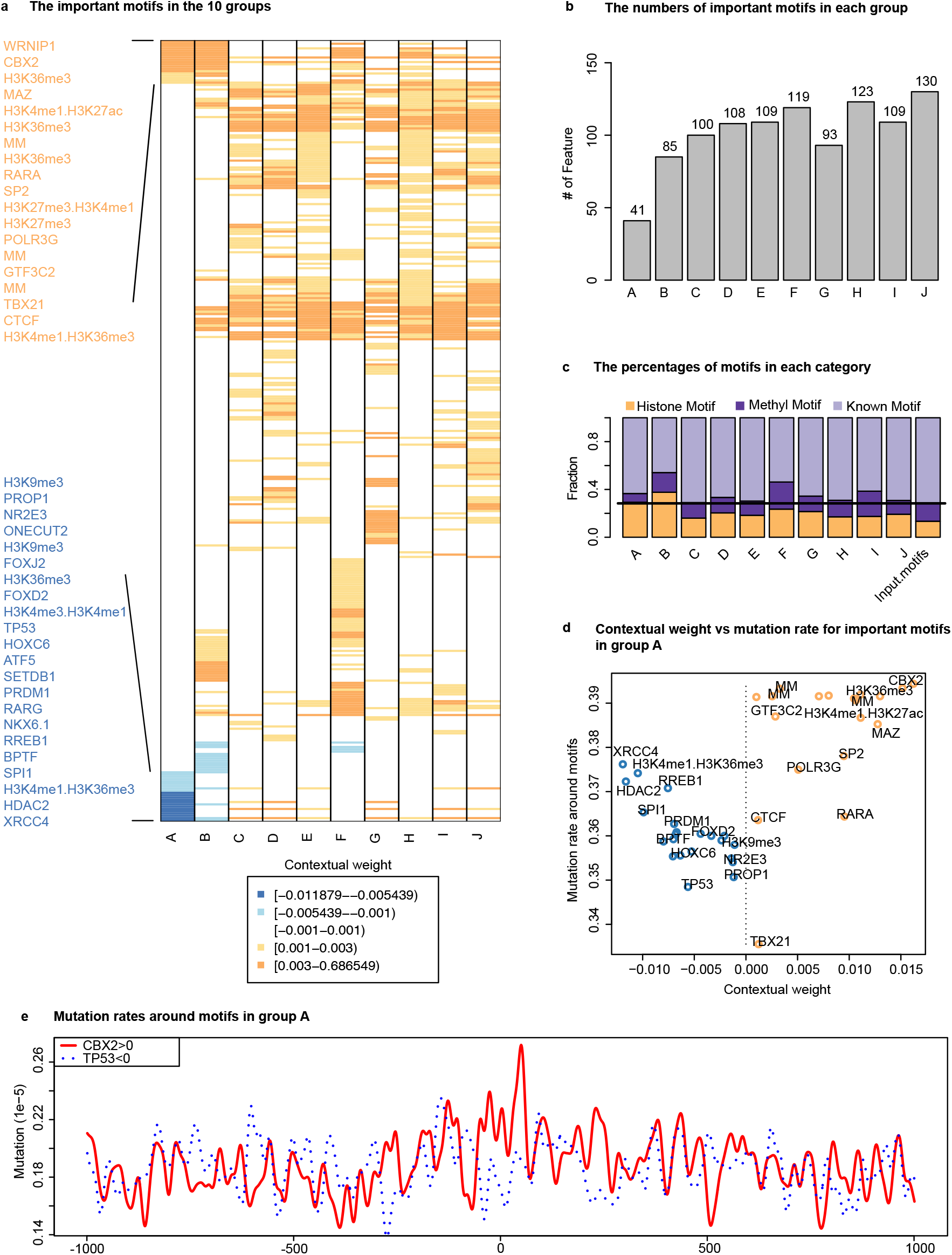
Identification of important motifs in cancer-independent regions. (**a**) The important features in the 10 groups. Blue and orange represent the negative and positive contextual weight, respectively; (**b**) The number of important motifs in each group; (**c**) The percentages of motif categories in each group; (**d**) The average mutation rates around the motifs with positive contextual weights are higher than those around the motifs with negative contextual weights in the group A regions; (**e**) Mutation rates around the motif sites (1kbp at each side of the motif) in the group A regions. The red and blue lines represent the motifs with positive and negative contextual weights, respectively. The motif site is at the center.

**Figure 5.**
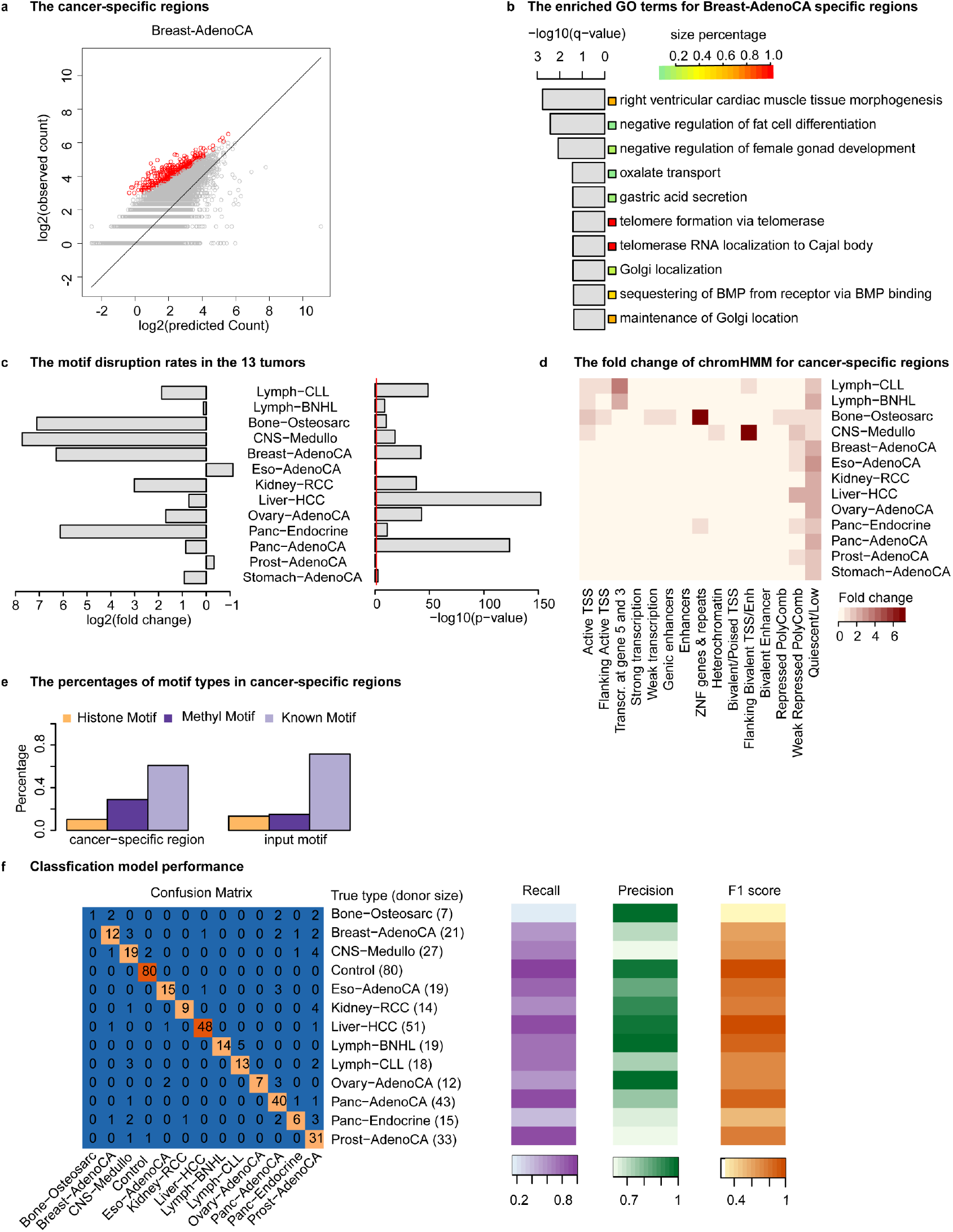
Analysis of the cancer-specific regions. (**a**) The identified cancer-specific regions in Breast-AdenoCA (red dots); (**b**)The enriched pathways for the cancer-specific regions in Breast-AdenoCA; (**c**) The fold change and p-value for the motif disruption rates in the 13 tumors. The red line represents the p-value of 0.05; (**d**) The fold change of chromHMM state (same as that in Figure 3 e) for the cancer-specific regions in each tumor; (**e**) The percentages of motif types that were significantly disrupted in cancer-specific regions; (**f**) The classification model performance. The confusion matrix for the classification model using the 150 selected cancer-specific regions on the testing dataset. Rows and columns correspond to the true and predicted tumor types, respectively. Values are the number of donors classified correctly. For example, for the 33 Prost−AdenoCA 33 donors, 31 donors were correctly classified.

Furthermore, we merged the similar motifs to remove redundancy and the model performed comparably with the original model (Supplementary Table 7). We chose to use the non-merged motif set in this study because different versions of the same motifs may represent collaborations of the same TF with different partners.

### Contextual regression identified important features in cancer-independent regions

To identify the most important features in each cancer-independent region, we selected motifs with both the largest contributions to the predicted value and the largest contextual weights: (1) we first selected the top 10% of features with the largest |β_i_X_i_|, where βi is the contextual weight for feature X_i_. |β_i_X_i_| represents the contribution of feature X_i_ to the predicted mutation rate of this region. (2) Among the top 10% of features with the largest |β_i_X_i_|, we selected the top 10% of features with the largest |β_i_|. In the following analyses, we only focused on the selected motifs and each region was thus represented by a vector composed of the contextual weight value βi for the selected motifs and 0 for the unselected motifs.

Using the contextual weight profile in each cancer, we clustered the cancer-independent regions into 10 clusters using K-means. After removing the clusters with small size (less than 10 regions), we obtained a total 123 clusters in 13 cancers and they were further clustered into 10 groups using hierarchical clustering with the most clusters of 16 in group A and the least of 4 in group G (Figure 3 b, see Supplementary Figure 6 for the full heatmap). The 10 groups show distinct characteristics. First, the mutation rate is the lowest in group A and the highest in group G (Figure 3 c). Second, the 10 groups have significantly different numbers of regions, with the largest number of 666,301 in group A and the smallest number of 331 in group G (Figure 3 d). Third, two ChromHMM states, the Weak Repressed PolyComb and Quiescent/Low states, were only enriched in group A across all 13 cancers but not in any other group (Figure 3 e and Supplementary Figure 7).

To get the important features in each group, we calculated the average of contextual weights across clusters for each group (Figure 4a and Supplementary Table 8). In total, 341 unique motifs were identified across the 10 groups, with the largest number of 130 in group J and the smallest number of 41 in group A (Figure 4 b). Interestingly, although epi-motifs (histone modification and DNA methylation associated motifs) only accounted for 28% of the input motifs, their percentage increased to average 36% in the important motifs in the 10 groups and this increase was particularly high in group A, B, F and I (Figure 4 c). This significant increase supports the roles of epi-motifs in establishing regional epigenetic states (22, 23).

We inspected group A as it has the most regions. We found 19 motifs with positive and 22 motifs with negative contextual weights in group A (Figure 4 a). The sign of the contextual weights indicates positive or negative association between the input feature and the predictive value. Therefore, our analysis suggested that the 19 and 22 motifs would have opposite impacts on somatic mutations. We checked the mutation rates around these motifs. Overall, the motifs with positive weights show significantly higher mutation rates than those with negative values (p-value of 1.06×10^−6^ from Student’s t-test) (Figure 4 d). Next, we performed pairwise comparison between all the possible positive- and negative-weighted motif pairs. The mutation rates around the motif sites (upstream 50bp to downstream 50bp) for positive-weighted motifs were higher than the paired negative motifs in 65.9% (494/750, p-value cutoff of 0.05 using Student’s t-test) of all the pairs. One motif pair of CBX2 (positive coefficient) and TP53 (negative coefficient) is shown as an example in (Figure 4e). The mutation rate is significantly higher (*p*-value of 5.05×10^−4^ from Student’s t-test) around the CBX2 motifs than that around TP53 (i.e. M6403_1.02) motifs in group A. TP53 plays a prominent role in repairing damaged DNA (70) and its negative association with mutation rate is reasonable.

Among the important motifs in group A, there are 7 positive- and 5 negative-weighted histone-motifs (Figure 4a). The positive association of the H3K27me3 motifs and the negative association of H3K4me3.H3K4me1 and H3K4me1.H3K36me3 motifs with the mutation rate are not surprising and consistent with the literature. The positive association of 1 H3K4me1.H3K27ac, 1 H3K4me1.H3K36me3 and 3 H3K36me3 as well as negative association of 2 H3K9me3 motifs suggest the relationship between histone modification and somatic mutation at the kilobase scale is more complex than expected from the previous megabase-scale studies. Interestingly, we also found 3 DNA methylation motifs (MM) positively associated with somatic mutation rates, indicating the possible roles of DNA methylation on affecting DNA damage and repair.

### Cancer-specific genomic regions and motifs

Using the predicted mutation rates, we identified cancer-specific regions (see Online Methods) and their percentages range from 0.2% in CNS-Medullo to 10.5% in Prost-AdenoCA, with a mean of 4.1% (Figure 3 a). The small portion of these regions confirmed that the majority of the genomic regions are cancer-independent. One example for cancer-specific regions in Breast-AdenoCA is shown in Figure 5a. We analyzed the biological processes in cancer-specific regions for each cancer type (Supplementary Table 9) and the enriched ones for Breast-AdenoCA are shown in Figure 5b, including those relevant to breast tissue and/or breast cancer such as “right ventricular cardiac muscle tissue morphogenesis”, “negative regulation of fat cell differentiation”, and “negative regulation of female gonad development” as well as “telomere formation via telomerase” and “telomerase RNA localization to Cajal body”. The key gene WRAP53 is also found in the Breast-AdenoCA-specific regions, which is related to the increased risk of breast cancer (71–73).

We also found that the disruption rate of motifs by somatic mutations in cancer-specific regions was significantly higher than that in cancer-independent regions for 10 out of 13 tumor types (p-value < 0.05, Figure 5c). The disruption rate of motifs for cancer-specific regions is defined as the number of all motif binding sites overlapped with somatic mutation divided by the number of all motifs binding sites and the number of somatic mutations (see Online Methods). This observation supports the hypothesis that the DNA motifs help to shape the epigenetic state and mutations disrupting the motifs may be associated with tumorigenesis.

To identify the most significantly disrupted motifs in the cancer-specific regions for each cancer, we selected motifs that had a disruption rate in the top 5% of all motifs with a p-value < 0.05 (p-value calculated from the Student’s t-test for the alternative hypothesis of the disruption rate in cancer-specific regions larger than that in cancer-independent regions). In total, 342 motifs were obtained in 13 cancer types and 60.8% of them were known motifs, decreased from 72% in the input motifs (Figure 5e). 102 (28.9%) DNA methylation associated motifs represented a significant increase from 13% in the input motifs. Notably, all of them are unmethylated motifs (i.e. UM motifs that are associated with low DNA methylation level (23)). This observation is consistent with that hypomethylation is often observed in both highly and moderately repeated DNA sequences including heterochromatic DNA repeats in cancer (74). The most common motifs found across cancer types are 1 known motif (E2F1, motif ID: M4536_1.02), 2 unmethylation motifs (UM_235.9_3.32_0.65_1_SGCWCGCGGCGGC and unmethylation motif UM_326.6_2.71_0.59_6_CGCGCCCCGY). E2F1 plays a crucial role in cell cycle (75) and DNA repair (76). Somatic mutations in E2F1 binding sites could result in dysfunction of E2F1. We also found many motifs only disrupted in specific cancer types (Supplementary Table 10), such as M2321_1.02 (TP63) in Bone-Osteosarc, M6446_1.02 (RARG) in Breast-AdenoCA, and M5371_1.02 (EGR4) in Lymph-CLL.

We noticed that the cancer-specific regions in most cancer types are enriched with the ChromHMM state of “Quiescent/Low” that have no or low epigenetic signals. Furthermore, each cancer type also has its own specific epigenetic state. For example, the state of “Transcription at gene 5’ and 3’” is the most enriched in Lymph-CLL and Lymph-BNHL, and “ZNF genes & repeats” in Bone-Osteosarc and Panc-Endocrine (Figure 5 d).

### The cancer-specific regions are predictive of cancer types

The cancer-specific regions are presumably important in specific cancers. Therefore, we examined whether these regions could be predictive of cancer types. Twelve cancers with WGS data with >30 donors each were used for multi-class classification of cancer types. Stomach-AdenoCA was excluded due to the small sample size. We also included 400 healthy donors from GTEX as controls. Using the mutation counts in the 53,223 cancer-specific regions (the coverage of cancer-specific regions in each cancer listed in Supplementary Table 6 and these regions combined to span the entire genome), we trained a Gradient Boosting Decision Tree to classify 12 tumor types and control. 80% of cancer donors (i.e. the dataset used in CR model construction) and controls were used to train and evaluate the classification model. The remaining samples (independent data that were not used in the CR model construction) were used as the testing set. We systematically searched the hyperparameter space (27,000 combinations of 6 parameters, Supplementary Table 11 and method section) using 5-fold cross validations on the training data (i.e. the 80% of the 1,382 cancer and 320 control donors). The best parameter combination was selected based on the minimal difference of accuracy between the training and validation datasets (i.e. no overfitting). The prediction accuracy on the test data in the 5-fold cross validations was 0.865. We next re-trained the classification model using all the 80% data with the best parameter combination and its prediction accuracy on the left-out 20% dataset was 0.858.

Next, we conducted feature selection using the feature importance metric of Gradient Boosting Decision Tree and selected 150 regions (accounting for 5.8% of the whole genome) to re-train the model. We also performed the parameter tunings using 5-fold cross validation on the 80% dataset. The accuracy on the testing samples was 0.813 using the best parameter combination (Supplementary Table 12). Next, the classification model was re-trained using the 80% dataset with the best parameter combination and its prediction accuracy on the left-out 20% donors was 0.822 (Figure 5f). The recall for each class on the testing dataset ranged from 0.14 in Bone-Osteosarc to 1.00 in control, with a median of 0.72. The precision ranged from 0.62 in Breast-AdenoCA to 1.00 in Bone-Osteosarc, Lymph-BNHL and Ovary-AdenoCA, with a median of 0.83. The F1 score, a comprehensive metric, ranged from 0.25 in Bone-Osteosarc to 0.98 in Control, with a median of 0.75. It is not surprising that the best performance was achieved on control samples because it is easier to distinguish normal from tumor samples than to discriminate one tumor from another tumor. The performance on Bone-Osteosarc was the worst, which was likely due to small sample size (only 28 donors in training and 7 in testing). Overall, the performance of the model is satisfactory considering only 5.8% of the genome was used.

## Discussion

Distinct from the previous analyses, we showed for the first time that DNA motifs can be predictive of regional somatic mutation rates at the kilobase scale. The DNA motifs include known TF motifs as well as epi-motifs that are associated with histone modifications or DNA methylation. We showed that the prediction performance is comparable to that using histone and TF ChIP-seq data. Considering that genome sequencing is much easier than ChIP-seq experiments in tumor tissues, our model provides a powerful approach to quantify the relationship between somatic mutations and epigenetic state.

Notably, the epi-motifs account for a significant portion (average 36% in the 10 groups) of the motifs most predictive for mutation rates despite that only 28% of the input motifs are epi-motifs. Particularly, among the motifs that are disrupted by somatic mutations in the cancer-specific regions, DNA methylation associated motifs counted for 28.9%, a big jump from its percentage (13%) in the input motifs. Similar to that TF binding is crucial for transcriptional regulation and thus gene expression can be predicted by TF motifs, the pivotal roles of epi-motifs in predicting regional somatic mutation rates and their more frequent disruption in the cancer-specific regions indicate that they are important in shaping the regional epigenetic state and deciding the locus-specificity of epigenetic modifications. The important epi-motifs identified from this analysis can guide future studies to identify the proteins whose binding to these epi-motifs could recruit epigenetic enzymes to specific loci and initiate local change of the epigenetic state. The success of this study also suggests that it is possible to use epi-motifs as the surrogate of local epigenetic state for predicting other observable measurements.

We hypothesized that the majority of somatic mutations are random and only depend on the regional epigenetic state that constrains the DNA damage and repair machineries. In other words, the majority of the genomic regions containing somatic mutations are cancer-independent. This hypothesis was supported by the high correlation between the predicted and measured mutation rates in all the regions. We found that this kilobase-scale relationship is general across cancers as indicated by the successful prediction on cancers not included in the training dataset. Such a relationship cannot be uncovered from the megabase analysis in the previous studies.

Furthermore, the contextual regression model provides a framework to interpret the neural network predictions and identify the most predictive features. Using the contextual weights of the predictive features, we could cluster the genomic regions that share similar motif contribution profiles on predicting mutation rates. The regions in the same cluster are presumably regulated by similar mechanisms, which is similar to genes sharing similar expression profiles across cell types. In fact, by analyzing these clusters, we observed that the impact of protein binding on regional mutations can be positive, negative or neutral. While the previous study reported that TF binding would block DNA repair proteins in the open chromatin regions to increase mutation rates (55), we found there exist proteins/motifs whose occurrence in the open chromatin regions is associated with lower mutation rates. This observation highlights the importance of genetic and epigenetic context on impacting regional mutations.

Importantly, the predicted mutation rates from the contextual regression model provide a quantitative background to identify cancer-specific regions in a particular cancer that have significantly higher mutation rates than expected from the regional epigenetic states in the corresponding normal tissue. The CR model learns the relationship between features and mutation rate. For each region from each cancer, it has its own background mutation rate based on its own features. Therefore, if the observed mutation rate is higher than the background, the region may be related to the mechanism of this cancer. In this study, we call it cancer-specific region. Some of these regions may be common in some extent since the different cancer may share some common characteristic. Some of these regions should be different because of different cancers. Based on these cancer-specific regions, we showed that only using 150 features/regions that account for only 5.8% of the human genome could predict cancer types with a satisfactory performance. This result provides a potential diagnosis tool using targeted sequencing.

In summary, several interesting observations can be drawn from this study. First, using epi-motifs to predict biological outputs depending on local epigenetic state such as somatic mutation rates would help to illustrate their importance in regulating epigenetic locus-specificity. The significant portion of epi-motifs among the important motifs we identified confirmed the association between epi-motifs and epigenetic state. Second, while the epigenetic state is known associated with regional mutation rate, no prediction model has been established at kilobase resolution and our model is the first to achieve this resolution. We showed that the majority of the genomic regions containing somatic mutations are cancer-independent and such a relationship cannot be uncovered from the megabase analysis in the previous studies. Third, the contextual regression allows identification of important motifs contributing to the regional mutation rates, which is also the first study to uncover motifs from computational models regulating mutation rates. Fourth, the predicted mutation rates from the contextual regression model provide a quantitative background to identify cancer-specific regions in a particular cancer, which can be used to classify cancer types and provide more insights of underlying mechanisms.

During the submission of this work, Sherman et al. (83) published a deep learning method to predict mutation rates at 10kbp resolution. Our study is different from Sherman et al. (83) in the following aspects and the two studies are highly complementary. First, our model only uses DNA motifs and epigenomic data of 6 histone marks in the corresponding normal cell types for each cancer type. In contrast, Sherman et al. (83) used much more data, including 723 chromatin marks from 111 tissues to train their model as well as replication timing in 10 cell lines and average nucleotide and CG content in the reference genome. Our model uses much less features but can achieve a slightly better prediction performance (a mean Pearson R2=0.736, i.e. R=0.858, Fig. 2B and Supplementary Table 3 in our manuscript compared to a mean Pearson R2=0.706, i.e. R=0.84, in Sherman et al. (83)). Second, our model can be applied to new cancer types without retraining and only requires the epigenetic state in the corresponding normal cells as shown in our paper that the model trained on 5 cancer types was successfully applied to predict the remaining 8 cancer types. In contrast, Sherman et al. (83) has to re-train the entire model to include new cancer types. Third, our study focuses on understanding how the DNA motifs particularly the epigenetic motifs contribute to regional somatic mutation rates and the contextual regression model provides an interpretable model that directly derives the important motifs. In contrast, Sherman et al. (83) focused on uncovering driver mutations.

## Materials and Methods

### Somatic mutation data

The somatic mutation data of 2,583 donors analyzed by the Pan-Cancer Analysis of Whole Genomes Consortium (PCAWG) were downloaded from the International Cancer Genome Consortium (ICGC) data portal (77). This was the largest data set when this analysis started. The tumor types for the donors were retrieved from Supplementary Table 1 in reference (68). We filtered the data using the following criteria: (1) donors with metastatic tumors were removed because in this study we focused on the primary tumors; (2) outlier donors with extremely high or low somatic mutation numbers were discarded to avoid bias. An outlier was defined as a data point located outside 1.5 times the interquartile range above the upper quartile and below the lower quartile for a tumor type; (3) tumor types with less than 5 donors or without ChromHMM segmentation in the corresponding normal tissues were not included; (4) only WGS data were kept if a donor had both WES and WGS data; (5) removing tumor types if the number of the total somatic mutations in all the donors of that tumor type was less than 30,000. As a result, 1,125 donors from 13 different tumor types remained for model training and testing (Supplementary Table 1). Somatic mutations in the blacklisted regions were removed (78).

### DNA Motifs

We included 1731 human motifs of DNA binding proteins documented in the CIS-BP database (79) and another 55 motifs from Factorbook (64). We also added 313 motifs associated with DNA methylation (23) and 361 motifs associated with histone modifications (22) that were identified in our previous studies. These motifs were used to approximate the epigenetic state. We used FIMO (80) to scan these total 2460 motifs against hg19. With a p-value cutoff of 10^−5^, 2,321 motifs having at least one occurrence were used for the following analyses (Supplementary Table 4).

### Genomic segmentation using ChromHMM

The core 15-state ChromHMM segmentations were downloaded from https://egg2.wustl.edu/roadmap/web_portal/. For kidney and prostate gland, the ChromHMM segmentations were not available from the website. To be consistent, we applied the core 15-state trained ChromHMM model to these 2 tissues, which was downloaded from https://egg2.wustl.edu/roadmap/data/byFileType/chromhmmSegmentations/ChmmModels/coreMarks/jointModel/final/model_15_coreMarks.txt. The data for these 2 tissues were downloaded from the ENCODE portal (Supplementary Table 13).

### Somatic mutation density and feature calculation

We calculated the somatic mutation density in a given set of cancer patients as the following. Let *R*_*i*_ denote the segmented region *i, i* = 1, …, *N, l*_*i*_ the length (bp) of *R*_*i*_, and *C*_*i*_ the number of somatic mutations in all the donors in region *i*. The somatic mutations were downloaded from PCAWG. The regional somatic mutation density *D*_*i*_ was computed as 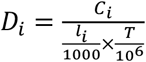, where *T* is the total number of somatic mutations in all the donors of this dataset. We added a pseudo-count to *D*_*i*_ and defined 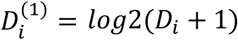. We then calculated 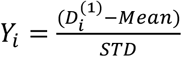 as the response variable in the model by z-score transformation, where *Mean* is the average of 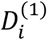 and *STD* is the standard deviation error of 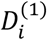. Let *M*_*j*_ be the motif *j*, 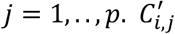 is the median p-value of all occurrences of *M*_*j*_ in the region *R*_*i*_ and the p-value was calculated from FIMO (80). We used 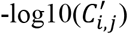 in each region as the input features to predict *Y*_*i*_.

### Training and testing of the CR model

The details of the training and testing of the CR model is shown in Supplementary Figure 1. Briefly, we divided the donors randomly into two sets:80% for model training/testing and the rest for independent testing. Using the 80% donors, we trained and tested CR models for individual cancer types using 10-fold cross validations (Step 1 in Supplementary Figure 1). The best performed 5 cancers were selected to train a universal model on all the regions in these 5 cancers (Step 2). As the majority of the regions in the cancers could be accurately predicted using the motifs and the ChromHMM segmentation in the corresponding normal cells, it confirmed our hypothesis that somatic mutations in the majority of the genome are cancer-independent. To better capture the relationship between the somatic mutations and epigenetic state, we removed the regions, i.e. cancer-dependent regions, whose predicted mutations rates significantly deviated from the observed values (Step 3). We further analyzed important motifs with high CR weights (Step 4). We then trained and tested a classification model to distinguish cancer types using the cancer-dependent regions on the 20% of the donors that were not used to select these regions as an independent test (Step 5).

### Assessing the CR model performance

In each dataset, we performed cross validation to assess the model performance, in which 10% of the segmented regions were held out for testing. Because there were overlapped ChromHMM regions from different tumor types, we partitioned the samples from these 5 tumor types based on the chromosomes to avoid the overlapped regions presenting in training and testing datasets. Two or three chromosomes were randomly left out for testing while the other chromosomes were used for training CR models. We repeated such cross validation 10 times. The specific partitions of training/testing dataset and performances were listed in Supplementary Table 5.

### Identification of cancer-specific and cancer-independent regions

To identify the cancer-specific and cancer-independent regions, we used an iterative procedure. First, we trained a CR model using all the regions from the merged dataset (the first iteration). As the majority of the regions are cancer-independent regions, using all the regions to train the model would not significantly impact the accuracy. Assuming the cancer-independent somatic mutation counts in a specific region follow a Poisson distribution, we estimated the parameter lambda (i.e. the expectation for Poisson distribution) using the predicted counts that were converted from the predicted mutation rates. Based on this Poisson distribution, we calculated a p-value for the observed mutation count. If a region had a p-value (upper-tail or lower-tail) < 0.1, it was considered as either cancer-specific or ambiguous and thus removed from the training set. We retrained the CR model using the remaining regions that presumably contained more cancer-independent ones in the second iteration. Repeating this procedure would continue improving the model and removing regions that are cancer-specific or ambiguous. We found that this procedure converged fast as the average mean squared error (MSE) on the testing dataset reached the plateau at the second iteration, indicating that the model became stable. Therefore, we took this model trained using two iterations as the final model for the following analyses.

We used the predicted mutation rates from the final CR model as the background, and re-calculated the p-value for each region in each tumor. A region was called as cancer-independent if its p-value (upper-tail and lower-tail) > 0.1, cancer-specific if the FDR from the upper-tail for a region was <0.01. The other regions were ambiguous regions and not included in any further analysis.

### Somatic mutation calling for the GTEX data

We downloaded the GTEX WGS data from dbGap (accession number phs000424.v8.p2). We aligned the sequencing reads to hg19 and called somatic mutations using the GATK best practice workflow (81). We removed the reads originated from duplicates of the same DNA fragments using MarkDuplicatesSpark function in GATK. Base (Quality Score) Recalibration was conducted for correcting any systematic bias observed in the base quality scores. We called candidate variants using Mutect2. FilterMutectCalls was applied to identify variants from artifacts, such as those resulted from alignment, strand and orientation bias, polymerase slippage, and germline variants. This generated a VCF file with a FILTER field. The true positives were labeled with PASS in the FILTER field. Funcotator was used to add annotation to these variants, such as dbSNP and gencode. Lastly, we only considered somatic mutations with a FILTER flag PASS and obtained somatic mutations for 400 donors.

### Disruption rate of the motifs

In the cancer-specific regions for a given patient, the disruption rate of a motif was calculated as C/(M*N), where C is the number of motif binding sites overlapped with somatic mutations in the cancer-specific regions in this patient, M is the total number of motif binding sites and N is the total number of somatic mutations in the cancer-specific regions for this patient. Similarly, we calculated the disruption rate in the cancer-independent regions. To test whether the disruption rate in the cancer-specific regions was higher than in the cancer-independent regions, paired-T test was used to compute the p-value for each tumor type with the patients as samples. This way, we identified significantly disrupted motifs for each cancer type.

To evaluate whether or not all motifs were significantly disrupted in one type of cancer, we performed the above analysis for all the motifs. Specifically, given the cancer type and a patient, the disruption rate for all motifs was defined as C/(M*N), where C is the number of all motif binding sites overlapped with somatic mutations in the cancer-specific regions in this patient, M is the total number of all motif binding sites and N is the total number of somatic mutations in the cancer-specific regions of this patient. The disruption rate in cancer-specific regions is calculated in the same way. Paired-T test was used to evaluate the significance for each tumor type with the patients as samples. The p-value cutoff was set to 0.05.

### Gradient Boosting Decision Tree

A gradient boosting decision tree was trained to classify cancer types using the scikit-learn package (82). There were six parameters in the model, including (1) learning rate (denoted as learning_rate); (2) the minimum number of samples (or observations) required in a node to be considered for splitting (min_samples_split); (3) the minimum samples (or observations) required in a terminal node or leaf (min_samples_leaf); (4) the maximum depth of a tree (max_depth); (5) the fraction of observations to be selected for each tree (subsample); (6) the number of sequential trees to be modeled (n_estimators).

We selected the optimal values of the parameters with the best classification performance: when using all the cancer-specific regions as features: learning_rate=0.012; min_samples_split=150; min_samples_leaf=130; max_depth=2; subsample=0.6; n_estimators=1900 (Supplementary Table 11); when using the selected 150 cancer-specific regions as features, learning_rate=0.011; min_samples_split=190; min_samples_leaf=60; max_depth=3; subsample=0.6; n_estimators=2000 (Supplementary Table 12).

## Data availability

The somatic mutation data was downloaded from the International Cancer Genome Consortium (ICGC) data portal (https://dcc.icgc.org/). The ChromHMM segmentations were downloaded from https://egg2.wustl.edu/roadmap/web_portal/. The Chip-seq data for GM12878 was downloaded from Encode (https://www.encodeproject.org/). The GTEX WGS data was downloaded from dbGap (accession number phs000424.v8.p2). The human motifs were from refs (22, 23, 64, 79).

## Acknowledgements

This work was partially supported by the NIH (R01HG009626 to W.W.).

## Author contributions

Z.W., J.W. and W.W. designed all the analyses. Z.W. and Cong L. performed all the analysis and prepared the figures, tables and contributed to the software development and its implementation. W.W. conceived and supervised the project. Chengyu L. contributed to the implementation of Contextual Regression. M.W. provided the methylation motifs and V.N. provided the histone motifs. Z.W., Cong L. and W.W. wrote the manuscript.

## References

1. S. H. Stricker, A. Köferle, S. Beck, From profiles to function in epigenomics. Nat. Rev. Genet. 18, 51–66 (2016).

2. K. Struhl, E. Segal, Determinants of nucleosome positioning. Nat. Struct. Mol. Biol. 20, 267–273 (2013).

3. J. Ernst, M. Kellis, Discovery and characterization of chromatin states for systematic annotation of the human genome. Nat. Biotechnol. 28, 817–825 (2010).

4. M. B. Stadler, et al., DNA-binding factors shape the mouse methylome at distal regulatory regions. Nature 480, 490–495 (2011).

5. J. L. Rinn, H. Y. Chang, Genome Regulation by Long Noncoding RNAs. Annu. Rev. Biochem. 81, 145–166 (2012).

6. A. I. Badeaux, Y. Shi, Emerging roles for chromatin as a signal integration and storage platform. Nat. Rev. Mol. Cell Biol. 14, 211–224 (2013).

7. J. R. Dixon, et al., Topological domains in mammalian genomes identified by analysis of chromatin interactions. Nature 485, 376–380 (2012).

8. E. P. Nora, et al., Spatial partitioning of the regulatory landscape of the X-inactivation centre. Nature 485, 381–385 (2012).

9. K. S. Zaret, J. S. Carroll, Pioneer transcription factors: establishing competence for gene expression. Genes Dev. 25, 2227–2241 (2011).

10. M. Levine, C. Cattoglio, R. Tjian, Looping Back to Leap Forward: Transcription Enters a New Era. Cell 157, 13–25 (2014).

11. A. Mayran, J. Drouin, Pioneer transcription factors shape the epigenetic landscape. J. Biol. Chem. 293, 13795–13804 (2018).

12. K. S. Zaret, Pioneer Transcription Factors Initiating Gene Network Changes. Annu. Rev. Genet. 54, 367–385 (2020).

13. G.-C. Yuan, Linking genome to epigenome. Wiley Interdiscip. Rev. Syst. Biol. Med. 4, 297–309 (2012).

14. E. M. Mendenhall, et al., GC-Rich Sequence Elements Recruit PRC2 in Mammalian ES Cells. PLOS Genet. 6, e1001244 (2010).

15. J. P. Thomson, et al., CpG islands influence chromatin structure via the CpG-binding protein Cfp1. Nature 464, 1082–1086 (2010).

16. C. A. Klattenhoff, et al., Braveheart, a Long Noncoding RNA Required for Cardiovascular Lineage Commitment. Cell 152, 570–583 (2013).

17. M. C. Tsai, et al., Long noncoding RNA as modular scaffold of histone modification complexes. Science 329, 689–693 (2010).

18. F. Baudat, et al., PRDM9 is a major determinant of meiotic recombination hotspots in humans and mice. Science 327, 836–840 (2010).

19. A. Bulut-Karslioglu, et al., A transcription factor–based mechanism for mouse heterochromatin formation. Nat. Struct. Mol. Biol. 19, 1023–1030 (2012).

20. Y. Costa, et al., NANOG-dependent function of TET1 and TET2 in establishment of pluripotency. Nature 495, 370–374 (2013).

21. J. W. Whitaker, Z. Chen, W. Wang, Predicting the human epigenome from DNA motifs. Nat. Methods 12, 265–272 (2015).

22. V. Ngo, et al., Epigenomic analysis reveals DNA motifs regulating histone modifications in human and mouse. Proc. Natl. Acad. Sci. U. S. A. 116, 3668–3677 (2019).

23. M. Wang, et al., Identification of DNA motifs that regulate DNA methylation. Nucleic Acids Res. 47, 6753–6768 (2019).

24. H. J. Bussemaker, H. Li, E. D. Siggia, Regulatory element detection using correlation with expression. Nat. Genet. 27, 167–171 (2001).

25. E. M. Conlon, X. S. Liu, J. D. Lieb, J. S. Liu, Integrating regulatory motif discovery and genome-wide expression analysis. Proc. Natl. Acad. Sci. U. S. A. 100, 3339–3344 (2003).

26. T. Lindahl, R. D. Wood, Quality control by DNA repair. Science 286, 1897–1905 (1999).

27. A. Sancar, L. A. Lindsey-Boltz, K. Ünsal-Kaçmaz, S. Linn, Molecular mechanisms of mammalian DNA repair and the DNA damage checkpoints. Annu. Rev. Biochem. 73, 39–85 (2004).

28. H. Shen, P. W. Laird, Interplay between the cancer genome and epigenome. Cell 153, 38–55 (2013).

29. A. Gonzalez-Perez, R. Sabarinathan, N. Lopez-Bigas, Local Determinants of the Mutational Landscape of the Human Genome. Cell 177, 101–114 (2019).

30. F. Supek, B. Lehner, Scales and mechanisms of somatic mutation rate variation across the human genome. DNA Repair (Amst). 81 (2019).

31. P. Mao, J. J. Wyrick, Organization of DNA damage, excision repair, and mutagenesis in chromatin: A genomic perspective. DNA Repair (Amst). 81 (2019).

32. K. D. Makova, R. C. Hardison, The effects of chromatin organization on variation in mutation rates in the genome. Nat. Rev. Genet. 16, 213–223 (2015).

33. A. Hodgkinson, Y. Chen, A. Eyre-Walker, The large-scale distribution of somatic mutations in cancer genomes. Hum. Mutat. 33, 136–143 (2012).

34. B. Schuster-Böckler, B. Lehner, Chromatin organization is a major influence on regional mutation rates in human cancer cells. Nature 488, 504–507 (2012).

35. Y. H. Woo, W. H. Li, DNA replication timing and selection shape the landscape of nucleotide variation in cancer genomes. Nat. Commun. 3 (2012).

36. L. Liu, S. De, F. Michor, DNA replication timing and higher-order nuclear organization determine single-nucleotide substitution patterns in cancer genomes. Nat. Commun. 4, 1–9 (2013).

37. P. Polak, et al., Cell-of-origin chromatin organization shapes the mutational landscape of cancer. Nature 518, 360–364 (2015).

38. M. A. M. Reijns, et al., Lagging-strand replication shapes the mutational landscape of the genome. Nature 518, 502–506 (2015).

39. F. Li, et al., The histone mark H3K36me3 regulates human DNA mismatch repair through its interaction with MutSα. Cell 153, 590–600 (2013).

40. S. X. Pfister, et al., SETD2-Dependent Histone H3K36 Trimethylation Is Required for Homologous Recombination Repair and Genome Stability. Cell Rep. 7, 2006–2018 (2014).

41. N. J. Haradhvala, et al., Mutational Strand Asymmetries in Cancer Genomes Reveal Mechanisms of DNA Damage and Repair. Cell 164, 538–549 (2016).

42. F. Supek, B. Lehner, Clustered Mutation Signatures Reveal that Error-Prone DNA Repair Targets Mutations to Active Genes. Cell 170, 534-547.e23 (2017).

43. S. Sasaki, et al., Chromatin-associated periodicity in genetic variation downstream of transcriptional start sites. Science 323, 401–404 (2009).

44. H. Ying, J. Epps, R. Williams, G. Huttley, Evidence that Localized Variation in Primate Sequence Divergence Arises from an Influence of Nucleosome Placement on DNA Repair. Mol. Biol. Evol. 27, 637–649 (2010).

45. M. Y. Tolstorukov, N. Volfovsky, R. M. Stephens, P. J. Park, Impact of chromatin structure on sequence variability in the human genome. Nat. Struct. Mol. Biol. 18, 510–516 (2011).

46. X. Chen, et al., Nucleosomes suppress spontaneous mutations base-specifically in eukaryotes. Science 335, 1235–1238 (2012).

47. S. Morganella, et al., The topography of mutational processes in breast cancer genomes. Nat. Commun. 7 (2016).

48. O. Pich, et al., Somatic and Germline Mutation Periodicity Follow the Orientation of the DNA Minor Groove around Nucleosomes. Cell 175, 1074-1087.e18 (2018).

49. A. J. Brown, P. Mao, M. J. Smerdon, J. J. Wyrick, S. A. Roberts, Nucleosome positions establish an extended mutation signature in melanoma. PLoS Genet. 14 (2018).

50. R. Katainen, et al., CTCF/cohesin-binding sites are frequently mutated in cancer. Nat. Genet. 47, 818–821 (2015).

51. Y. A. Guo, et al., Mutation hotspots at CTCF binding sites coupled to chromosomal instability in gastrointestinal cancers. Nat. Commun. 9 (2018).

52. P. Mao, et al., ETS transcription factors induce a unique UV damage signature that drives recurrent mutagenesis in melanoma. Nat. Commun. 9 (2018).

53. K. Elliott, et al., Elevated pyrimidine dimer formation at distinct genomic bases underlies promoter mutation hotspots in UV-exposed cancers. PLoS Genet. 14 (2018).

54. D. Perera, et al., Differential DNA repair underlies mutation hotspots at active promoters in cancer genomes. Nature 532, 259–263 (2016).

55. R. Sabarinathan, L. Mularoni, J. Deu-Pons, A. Gonzalez-Perez, N. Lopez-Bigas, Nucleotide excision repair is impaired by binding of transcription factors to DNA. Nature 532, 264–267 (2016).

56. J. Hu, O. Adebali, S. Adar, A. Sancar, Dynamic maps of UV damage formation and repair for the human genome. Proc. Natl. Acad. Sci. U. S. A. 114, 6758–6763 (2017).

57. M. B. Burns, N. A. Temiz, R. S. Harris, Evidence for APOBEC3B mutagenesis in multiple human cancers. Nat. Genet. 45, 977–983 (2013).

58. S. A. Roberts, et al., An APOBEC cytidine deaminase mutagenesis pattern is widespread in human cancers. Nat. Genet. 45, 970–976 (2013).

59. L. B. Alexandrov, et al., Signatures of mutational processes in human cancer. Nature 500, 415–421 (2013).

60. V. Aggarwala, B. F. Voight, An expanded sequence context model broadly explains variability in polymorphism levels across the human genome. Nat. Genet. 48, 349–355 (2016).

61. S. Nik-Zainal, et al., Landscape of somatic mutations in 560 breast cancer whole-genome sequences. Nature 534, 47–54 (2016).

62. P. Polak, et al., A mutational signature reveals alterations underlying deficient homologous recombination repair in breast cancer. Nat. Genet. 49, 1476–1486 (2017).

63. E. D. Pleasance, et al., A comprehensive catalogue of somatic mutations from a human cancer genome. Nature 463, 191–196 (2010).

64. J. Wang, et al., Sequence features and chromatin structure around the genomic regions bound by 119 human transcription factors. Genome Res. 22, 1798–812 (2012).

65. ENCODE Project Consortium, An integrated encyclopedia of DNA elements in the human genome. Nature 489, 57–74 (2012).

66. C. Liu, W. Wang, Contextual Regression: An Accurate and Conveniently Interpretable Nonlinear Model for Mining Discovery from Scientific Data (2017) (October 29, 2019).

67. C. Liu, Y.-C. Liu, H.-D. Huang, W. Wang, Biogenesis mechanisms of circular RNA can be categorized through feature extraction of a machine learning model. Bioinformatics 35, 4867–4870 (2019).

68. P. J. Campbell, et al., Pan-cancer analysis of whole genomes. Nature 578, 82–93 (2020).

69. J. Ernst, M. Kellis, ChromHMM: Automating chromatin-state discovery and characterization. Nat. Methods 9, 215–216 (2012).

70. A. B. Williams, B. Schumacher, p53 in the DNA-damage-repair process. Cold Spring Harb. Perspect. Med. 6 (2016).

71. L. Silwal-Pandit, et al., The Sub-Cellular Localization of WRAP53 Has Prognostic Impact in Breast Cancer. PLoS One 10, e0139965 (2015).

72. N. Pouladi, S. Abdolahi, D. Farajzadeh, M. A. H. Feizi, Haplotype and linkage disequilibrium of TP53-WRAP53 locus in Iranian-Azeri women with breast cancer. PLoS One 14, e0220727 (2019).

73. S. Mahmoudi, S. Henriksson, L. Farnebo, K. Roberg, M. Farnebo, WRAP53 promotes cancer cell survival and is a potential target for cancer therapy. Cell Death Dis. 2, e114–e114 (2011).

74. M. Ehrlich, DNA methylation in cancer: Too much, but also too little. Oncogene 21, 5400–5413 (2002).

75. S. Y. Tsai, et al., Mouse development with a single E2F activator. Nature 454, 1137–1141 (2008).

76. E. H. Choi, K. P. Kim, E2F1 facilitates DNA break repair by localizing to break sites and enhancing the expression of homologous recombination factors. Exp. Mol. Med. 51, 1–12 (2019).

77. T. J. Hudson, et al., International network of cancer genome projects. Nature 464, 993–998 (2010).

78. H. M. Amemiya, A. Kundaje, A. P. Boyle, The ENCODE Blacklist: Identification of Problematic Regions of the Genome. Sci. Rep. 9, 1–5 (2019).

79. M. T. Weirauch, et al., Determination and Inference of Eukaryotic Transcription Factor Sequence Specificity. Cell 158, 1431–1443 (2014).

80. C. E. Grant, T. L. Bailey, W. S. Noble, FIMO: scanning for occurrences of a given motif. Bioinformatics 27, 1017–1018 (2011).

81. G. A. Van der Auwera, et al., From fastQ data to high-confidence variant calls: The genome analysis toolkit best practices pipeline. Curr. Protoc. Bioinforma. 43, 11.10.1-11.10.33 (2013).

82. F. Pedregosa, et al., Scikit-learn: Machine Learning in Python. J. Mach. Learn. Res. 12, 2825–2830 (2011).

83. M. A. Sherman, et al., Genome-wide mapping of somatic mutation rates uncovers the drivers of cancer. Nat. Biotechnol. (2022).

