## SupplementaryMaterials for "Predicting regional somatic mutation rates using DNA motifs"

### Supplementary Table Legend

**Supplementary Table 1**. The tumor types and donor size analyzed in this study.

**Supplementary Table 2**. The ChromHMM state similarity between cancer and corresponding normal cell lines. Four cancer-normal cell comparisons are shown in terms of length percentage of similar states ("length.similar"). To be more specific, TssA and TssAFlnk are deemed as similar; TxFlnk, Tx and TxWk are similar; ZNF/Rpts, Het, ReprPC, ReprPCWk and Quies are similar.

**Supplementary Table 3**. The summary statistics for 13 tumor types in this study. "Correlation in Train set": the average of Pearson correlations of CR model on the training dataset across 10-fold cross validation. "Correlation in Test set": the average of Pearson correlations of CR model on the testing dataset across 10-fold cross validation. "Merge data to train CR": whether or not to use this tumor type data to train a unified CR model. "Correlation in final model": the Pearson correlation between true value and the prediction from the final CR model applied to all regions in a tumor. "number of regions after removing overlapped regions": the number of regions after removing the overlapped regions with the 5 merged datasets. "Correlation with no overlapped regions": the Pearson correlation between the true value and the prediction from the unified CR model applied to regions without overlaps in a tumor.

**Supplementary Table 4**. The motif ID and the corresponding proteins used in this study. The number of binding sites in each chromosome is also listed. "TotalNum" represents the total number of binding sites in the whole genome.

**Supplementary Table 5**. The 10-fold cross validation results. "chr in test": the chromosome ID used as the testing dataset. "MSE_test": mean squared error in the testing dataset. "Cor_test": the Pearson correlation in the testing dataset. “MSE_train”: the mean squared error in the training dataset. “Cor_train”: the Pearson correlation in the training dataset.

**Supplementary Table 6.** The chromHMM dataset used in this study and the number of cancer-specific regions for each cancer.

**Supplementary Table 7**. Training and testing results using merged motif as features. (The annotation of column names can be found in Fig. 2b legend).

**Supplementary Table 8**. The important features in each group. “GroupA_beta”: the coefficient of feature in group A. 0 represents that the feature is not important in the group. “GroupA1_mean”: the average of mutation rate in the regions with the corresponding motif binding sites in group A.

**Supplementary Table 9**. The enriched pathways for the cancer-specific regions for 13 tumor types.

**Supplementary Table 10**. The motifs significantly disrupted by the somatic mutations. For example, UM_3582.2_3.88_0.56_57_known. TEAD2 is significantly disrupted in three datasets: Lymph-CLL, Kidney-RCC, Ovary-AdenoCA.

**Supplementary Table 11**. The classification performance using all the cancer-specific regions as features.

**Supplementary Table 12**. The classification performance using selected the 150 most important regions.

**Supplementary Table 13**. The accession number for the histone modification used in this study to perform the ChromHMM segmentation.

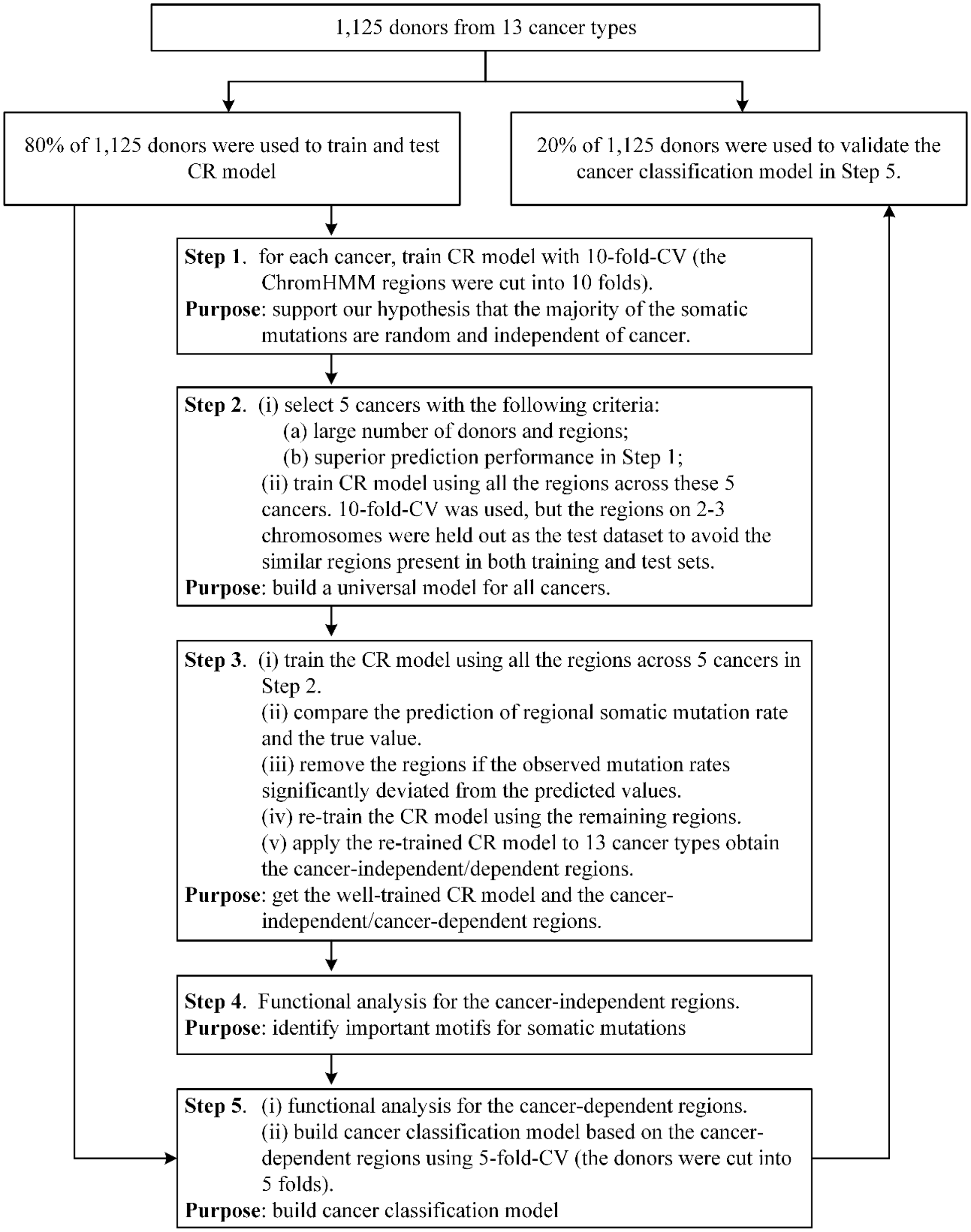

Supplementary Figure 1. The framework for the training and testing of the CR model.

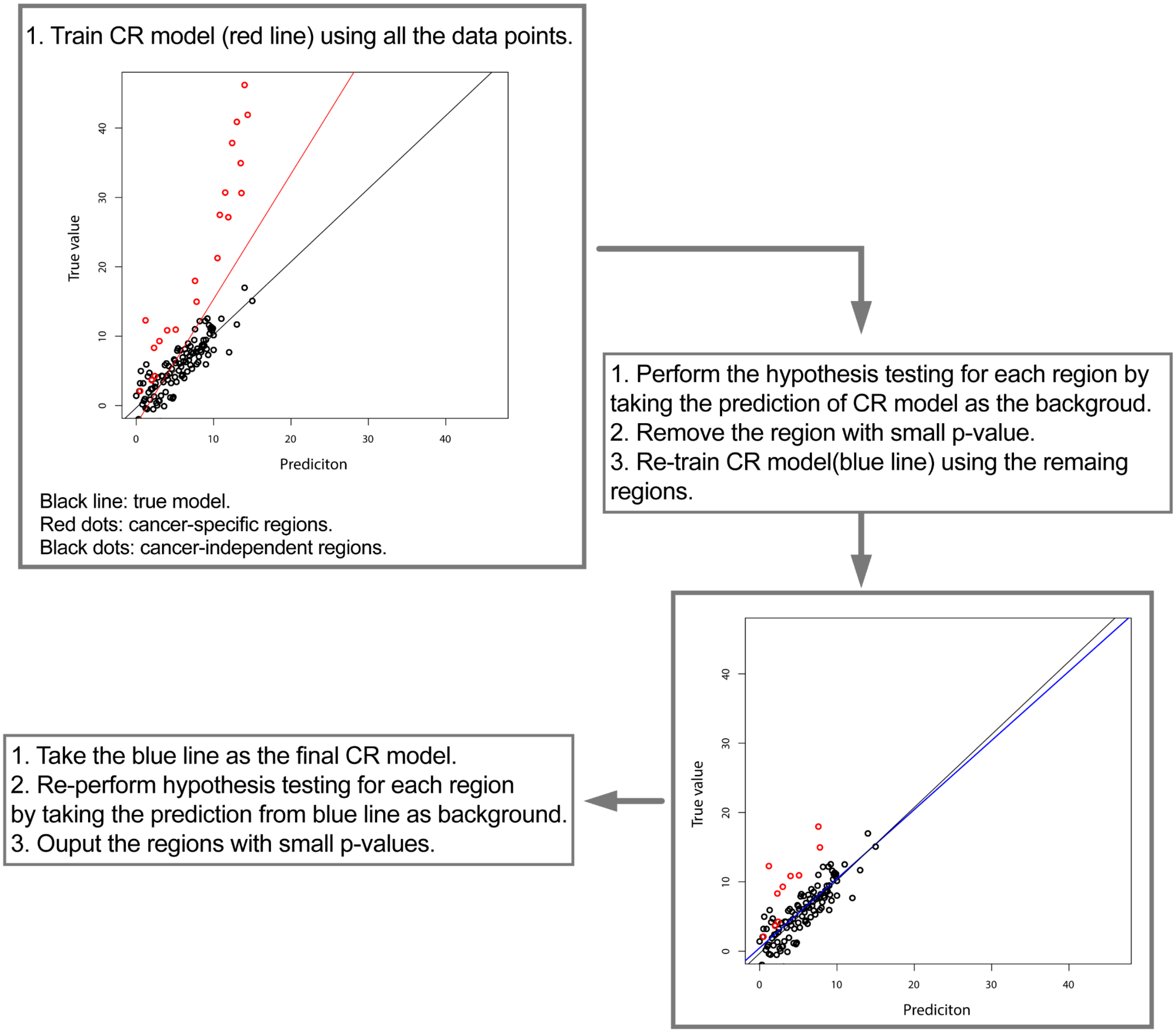


**Supplementary Figure 2**. The toy model illustrates how to train the CR model and identification of cancer-independent and cancer-specific regions. The red dots represent the cancer-specific regions and the black dots represent the cancer-independent regions. And the black line represents the true model. In practice, we don't know which dots (i.e. regions/samples) are cancer-specific or cancer-independent regions. So we train a CR model using all the samples and get the trained model indicated by the red line. Under the assumption, we know that the trained CR model is not exactly the true model, but it is close to the true model. To identify the cancer-specific regions, we take the prediction of the current trained CR model as the background and perform the hypothesis testing for each region (see Online methods for details). We remove the regions with small p-value and re-train the CR model using the rest of regions. And then a new CR model (i.e. the blue line) is obtained. This new CR model is closer to the true model. The blue line is treated as the true model and the hypothesis testing is done for each region again based on the prediction of the blue model. At last, the regions with small p-value will be taken as cancer-specific regions and regions with large p-value will be taken as cancer-independent regions.

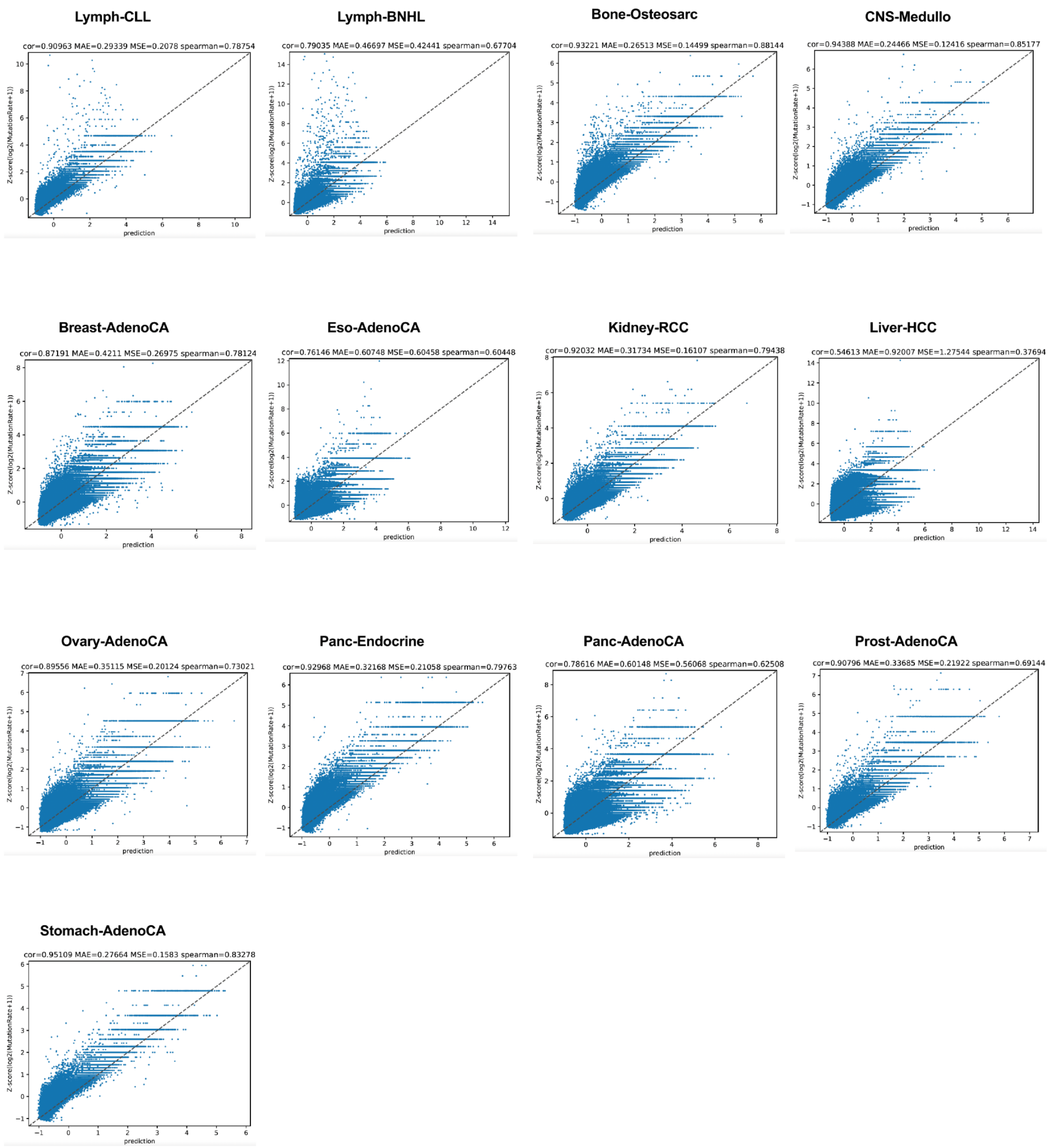


**Supplementary Figure 3.** The scatter plots for the 13 tumor types using the re-trained CR model. Cor: the Pearson correlation. MAE: mean absolute error. MSE: mean squared error. Spearman: Spearman correlation.

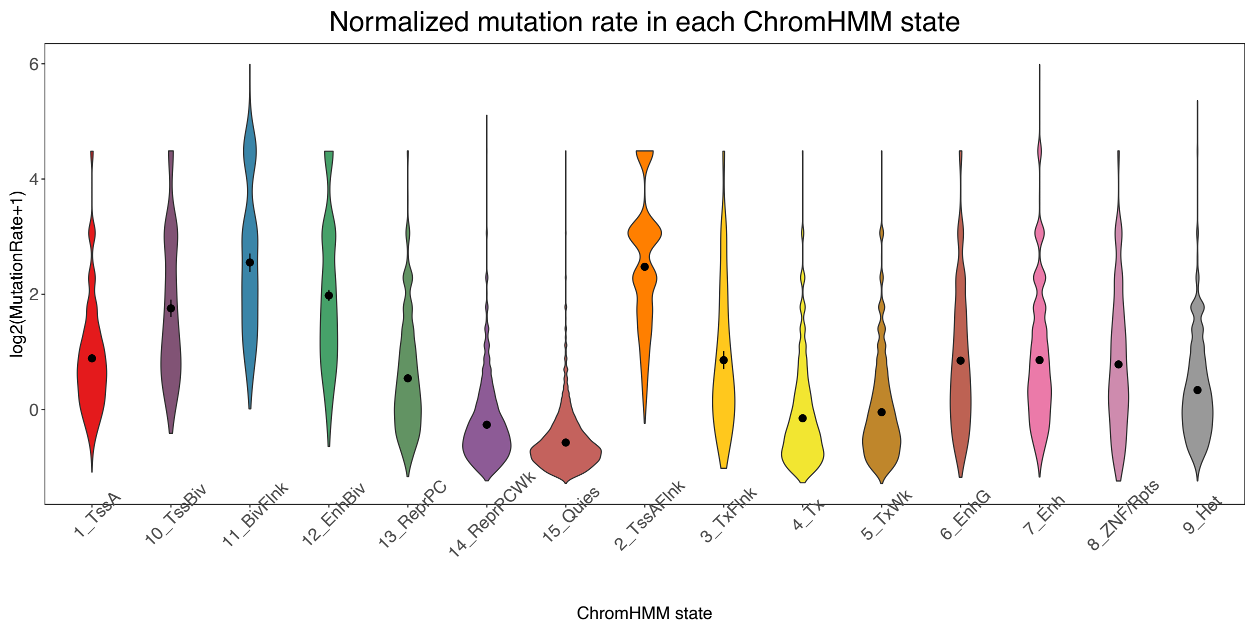


**Supplementary Figure 4**. The log2(MutationRate+1) distribution across ChromHMM states in breast cancer. All the other cancers have the similar wide distributions.

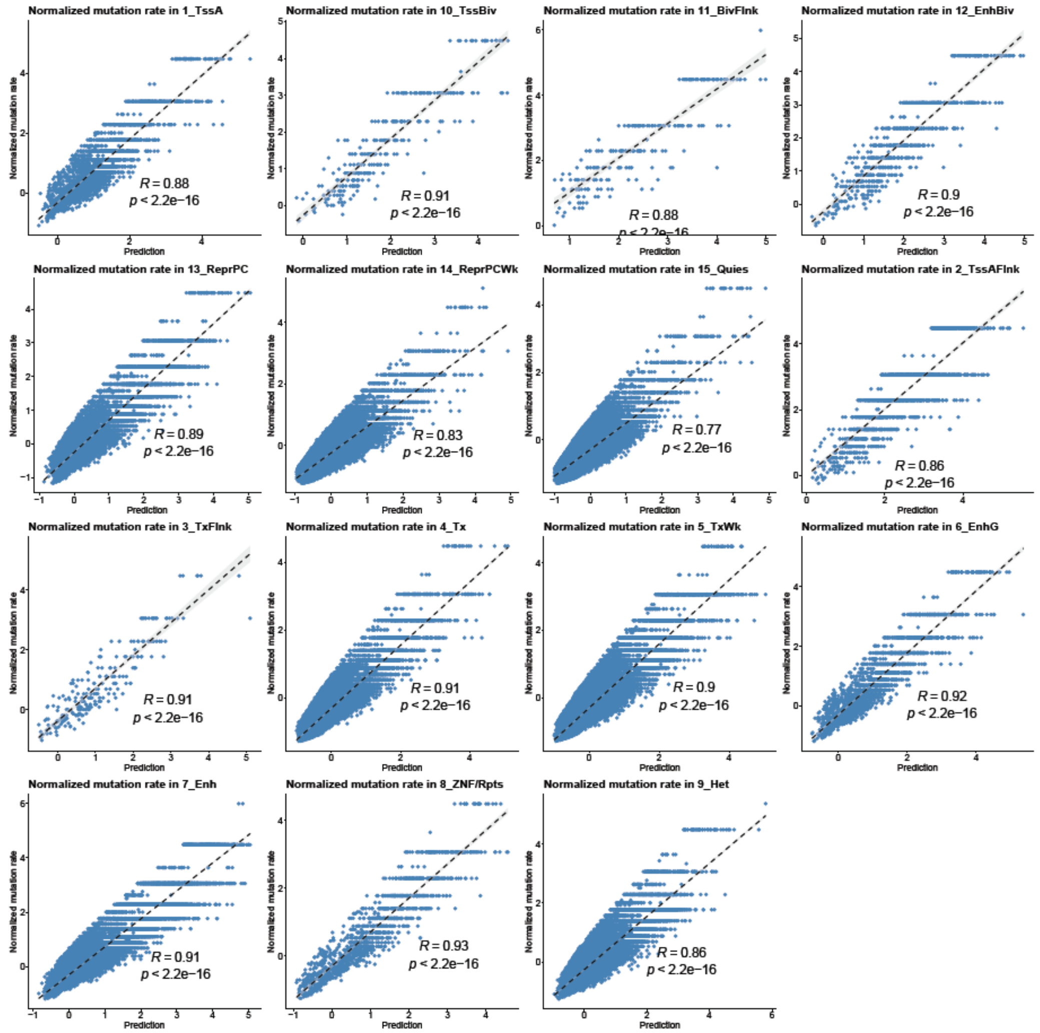


**Supplementary Figure 5**. Correlation between predicted and measured mutation rates across ChromHMM states in breast cancer. All the other cancers have the similar high correlations.

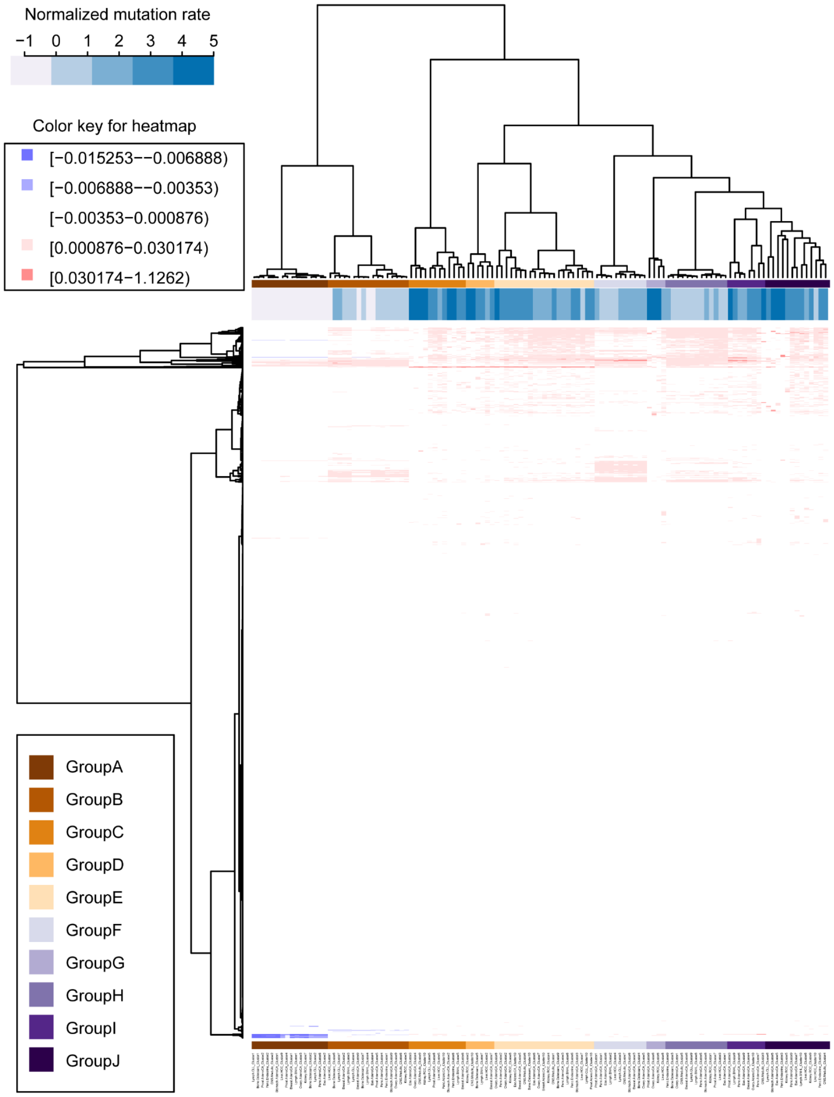


**Supplementary Figure 6.** Cancer-independent regions clustered using the contextual weights of the motifs. For each of the 13 cancer types, the identified cancer-independent regions were clustered into 10 clusters using the Manhattan distance between the feature contextual weight vectors as the similarity metric. Each row is a motif, each column a cluster, and each entry is the average of a motif's contextual weights in all the regions in a cluster. The clusters were further clustered into 10 groups.

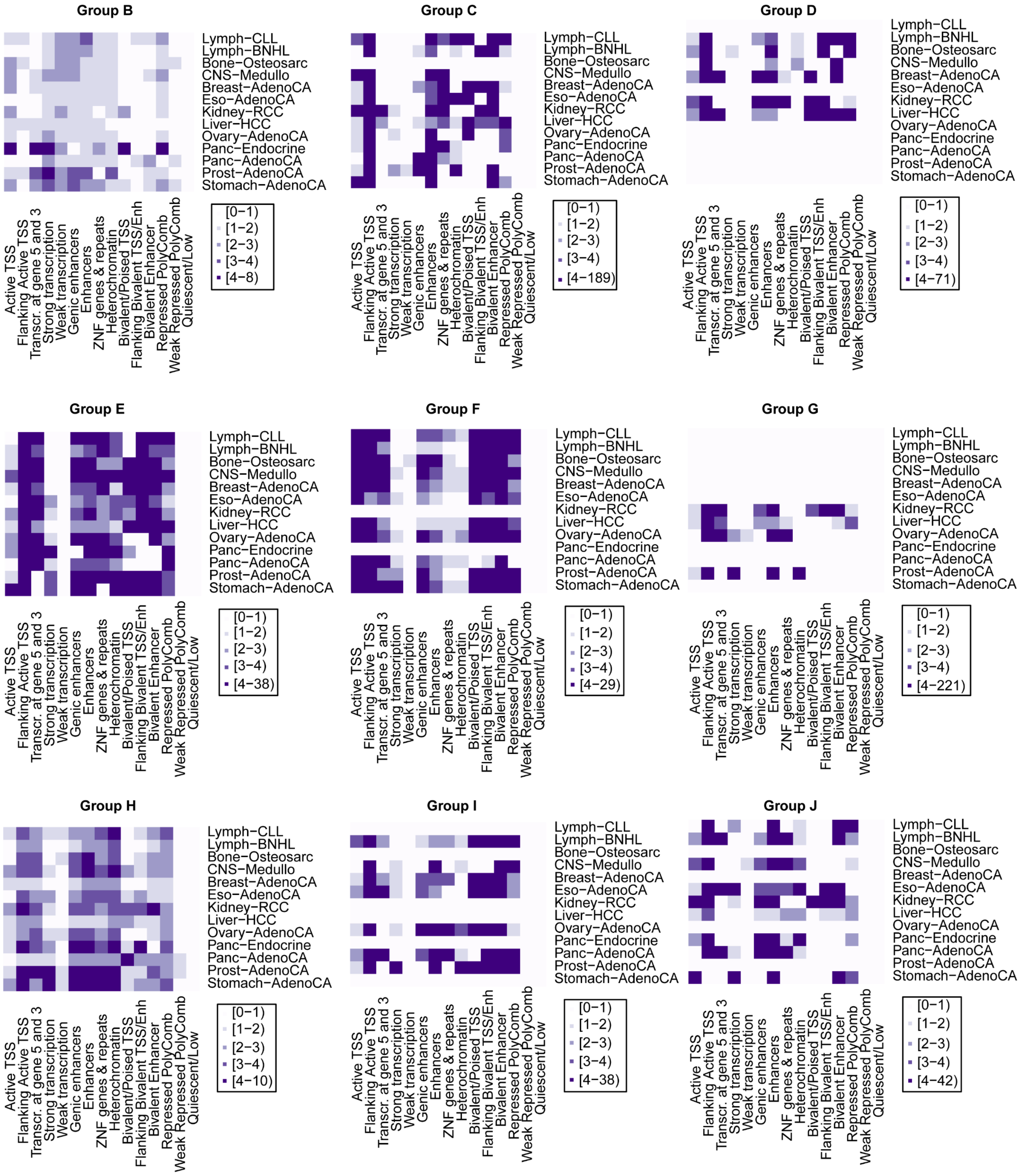


**Supplementary Figure 7**. The fold change of chromHMM state in the 9 groups from group B to group J. The color key represents the fold change between the percentage of one state in one group and the percentage of the state in the whole dataset.
